# Fright and Fight: Role of predation and competition on mate search tactics of wild male zebrafish

**DOI:** 10.1101/2020.03.28.013094

**Authors:** Aditya Ghoshal, Anuradha Bhat

## Abstract

Mate search tactics and association preferences among organisms in natural habitats can be dynamic and are determined by inherent trait preferences as well as the cost-benefit trade-offs associated with each mating decision. Two of the prime factors regulating mating decisions are the presence of competing conspecifics and predatory threats, both of which have important fitness consequences for the individual. We studied the influence of these two factors separately in mate search tactics and association preferences among zebrafish males. Male zebrafish were presented with a choice of two patches, consisting of different number of females, of which one patch was also associated with a predatory threat. We found that males made a preferential choice for the patch with more number of females only when the numerical difference between choices are starkly different, irrespective of the predatory threat associated with the patch. This points towards the role of numerical cognition in assessing cost-benefit tradeoffs in male zebrafish. We also studied the association preference of males in a multi-choice setup, consisting of four separate mixed-sex groups of zebrafish varying in densities. Our results showed that while test males preferred to visit the male-biased patches more often, they spent more time near female-biased patches or patches with equal sex ratio patches indicating the role of complex interplay of social cues in determining the associative behavior of males to a patch. This study, thus, sheds further light on the interactive roles of social cues and cognitive abilities in mate association patterns in this species.

## Introduction

Mating strategies in species, are driven by both innate preferences of individuals as well as several external environmental factors (Shuster and Wade 2003; Miles et al. 2007). Predation threat is one of the major ecological factors with fitness consequences influencing decision making during mate search and mate choice (Godin 1995; Barbosa and Magurran 2006). Immediate presence of predation threat requires the individuals to weigh the costs associated with mate search and mate sampling against the benefits. Similarly, presence of other competing individuals can increase the force of intra-sexual selection on males and might require a shift in their mating tactics. Presence of competitors can lead to aggressive interactions between males, reducing the overall fitness gains acquired from mating (Knell 2009). This study investigates the influence of these two major ecological factors on the mating strategies among wild zebrafish (*Danio rerio*).

Predation threat is one of the most important selective forces acting on an individual with direct lifetime fitness consequences (Lima and Dill 1990). Predation risk increases especially when the animal is searching for or displaying towards potential mates (Burk 1982; Endler 1992). Bright conspicuous male displays in many species also increase their predation risk (Zuk and Kolluru 1998). Thus, animals are under selection force to modulate their behaviors especially mate choice and mate search behaviors based on immediate predation risk (Lima and Dill 1990; Magnhagen 1991). It has been postulated that predation threat reduces the force of sexual selection (Forsgren 1992; Godin and Briggs 1996). Alternative mating strategies like sneaky matings increase in guppies in presence of an immediate threat (Godin 1995). In three-spined sticklebacks, males reduce their display behaviors when faced with an immediate predatory threat (Candolin 1997). In pipefish, males tend to have a preference for larger females but this preference is not maintained when the males sense a predation threat, upon which they tend to mate at random (Berglund 1994).

Operational sex ratio or OSR, defined as the ratio of reproductively receptive males to receptive females in a population (Emlen 1976), determines which sex will be under selection pressure to compete for access to mates (Kvarnemo and Ahnesjo 1996). How individuals modulate their mating tactics in response to competing individuals depend on the costs incurred during competition for access to the mates (Fawcett and Johnstone 2003). Competition increases the strength of sexual selection on the individuals which are in greater number and also make the prospective mates choosier, further increasing the strength of selection on the competing sex (Servedio and Lande 2006; Kokko and Johnstone 2002). There are several theoretical studies exploring the dynamics of male mate search and mating efforts with increased sperm competition (Weir et al. 2011; Fawceet and Johnstone 2003; Parker et al. 1996). Males show flexibility in mating-associated behavioral phenotypes in male-biased OSR (Weir et al. 2011). For example, male mice are found to modulate mating effort based on competition and low pay-off during sequential search for females (Ramm and Stockley 2014). Along with predatory threat and competition, mating tactics in males can also depend on the quality, receptivity and the number of females they encounter. Presence of higher density and abundance of females increases the chances of finding a suitable and receptive mate. This study investigates the response of male zebrafish to variation in female numbers in the presence of either predation or competition among a wild-caught population of zebrafish.

We examined the role of predation risk on association choice among male zebrafish using a two-choice test approach. This study examined the choice among males under varying female numbers and under predation risk. We performed a series of three experiments to assess male choice across patches with a gradient in female numbers, under different predation risk scenarios. To study the role of competition in association preferences among males, we performed multi-choice tests, with each patch varying in male: female sex-ratios. Specifically, a choice of four patches differing in male-female sex ratios (female-biased (1:3), no bias (1:1), male-skewed (1.67:1) and male-biased (3:1)) were provided to assess male preferences. We expected males to show preferential association with the female biased patches compared to male-skewed or male-biased patches, when we controlled for total number of individuals in the patches.

## Materials and Methods

### Procuring subject animals and maintenance

Wild-caught zebrafish (collected from Howrah district, West Bengal, India) were bought from a commercial supplier and maintained in the laboratory under 12L:12D conditions to mimic the natural LD cycle of zebrafish and temperature range of 23°C-25°C. The fish were kept in mixed sex-groups of approximately 60 individuals in well aerated holding tanks (60×30×30□cm) filled with filtered water and provided with standard corner filters for circulation. They were fed commercially purchased frozen dried blood worms once a day alternating with brine shrimp *Artemia spp.* The fish were maintained in the laboratory for six months before experiments were conducted to ensure they were all adults and were reproductively mature.

The study complied with the existing rules and guidelines outlined by the Committee for the Purpose of Control and Supervision of Experiments on Animals (CPCSEA), Government of India, the Institutional Animal Ethics Committee’s (IAEC) and guidelines of Indian Institute of Science Education and Research (IISER) Kolkata. All experimental protocols followed here have been approved by the Institutional Animal Ethics Committee’s (IAEC) and guidelines of Indian Institute of Science Education and Research (IISER) Kolkata, Government of India. No animals were euthanized or sacrificed during any part of the study, and behavioral observations were conducted without any chemical treatment on the individuals. At the end of the experiments, all fish were returned to stock tanks and continued to be maintained in the laboratory.

### Experimental setup for predation experiment

For the predation experiment we setup a standard choice test consisting of two identical chambers, one with a predator and one without (Figure 1). The test tank (dimensions: 60×30×30cms) consisted of a central open area with two chambers on either ends (dimensions of the chamber: 15×15×20cms) separated by a mesh partition (Figure 1). Each chamber was further divided into two sections where one of the chambers housed a predator fish (*Channa spp*.) on one section and stimuli females. This chamber represented the ‘predator-associated’ patch. At the chamber placed on the other end of the test tank, stimuli females were placed in one of the sections while the other section was kept empty. This chamber represented the ‘no predator’ patch. The center of the tank was provided with a removable holding-chamber with perforations to act as an acclimation chamber for the test males. The association preferences of test males were measured under three conditions that differed in terms of female numbers and presence/absence of predator. We varied the female number associated with the predator and the no-predator patches in the following manner:

**Figure 1.**
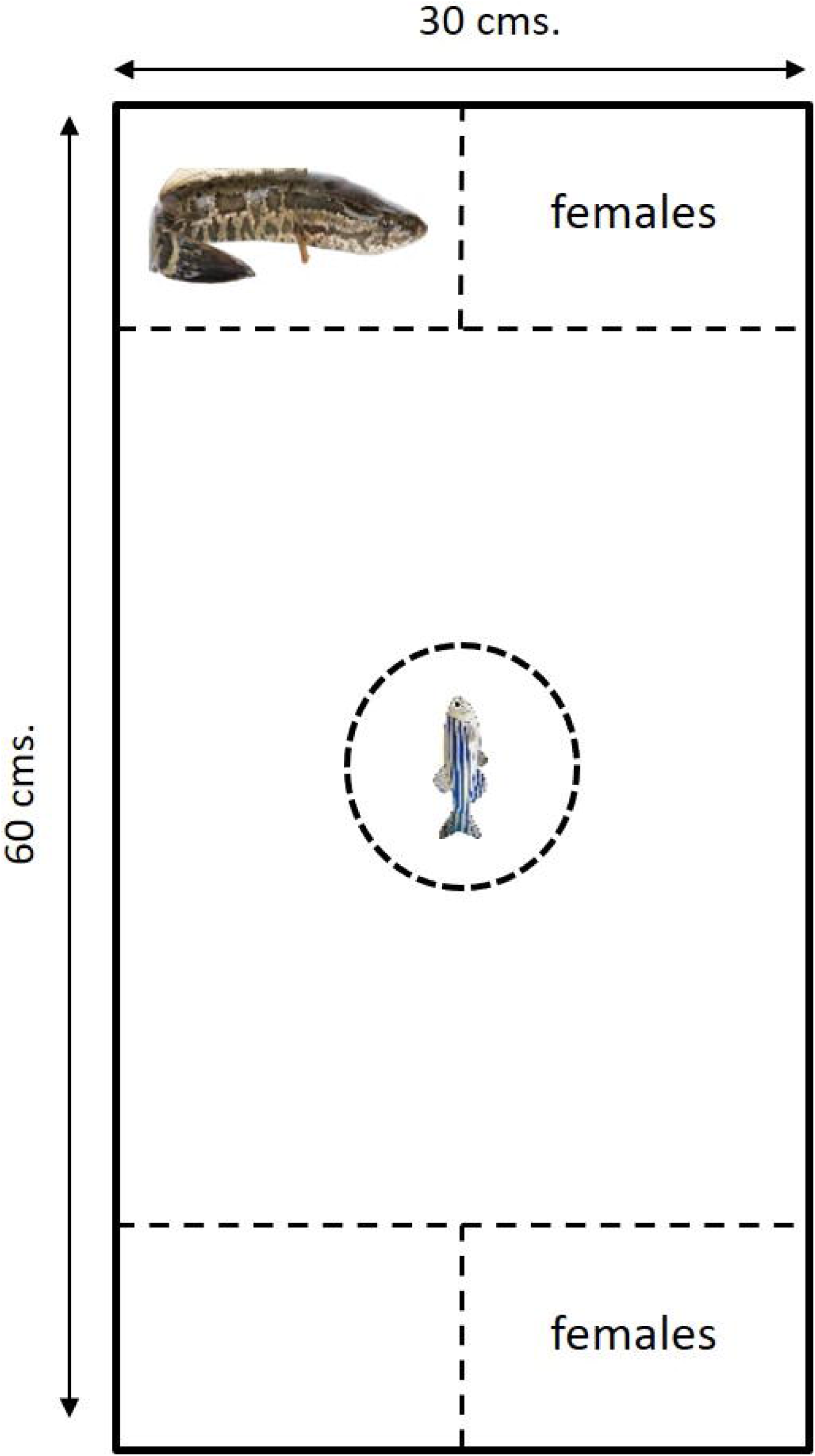
Schematic diagram of the experimental setup for testing male association preferences under predation threat. The central chamber contained the test males, in a transparent acclimation chamber with pores. This was removed remotely when the trials commenced, to release the male into the tank. A live predator *Channa spp.* is placed on side of the tank (predator side) and the other side is left empty (i.e. ‘no-predator’ side). Both predator and no-predator sides housed females separated from the predator/blank compartment by transparent mesh partitions.

E1 set: eight females with predator vs. two females without predator

E2 set: eight females with predator vs. four females without predator

E3 set: two females with predator vs. eight females without predator

We used the snakehead fish (*Channa spp.)*, a common predator and often found co-existing with zebrafish in natural habitats in the region (Spence et al. 2007). The predator fish were procured from a local fish supplier and housed in a holding tank in the laboratory under the same light and temperature conditions as the wild zebrafish population, for a month prior to experimentation.

We isolated test males four days prior to start of experiments and kept them in individual beakers with 300 ml filtered water. At the start of experimental observation, an individual test male was released in the acclimation chamber (a cylindrical plastic container, open at both ends, and with perforations) and left undisturbed for 7-10 minutes. Then the acclimation chamber was removed, allowing the test fish to move freely in the arena. The behavior of the test male was recorded for the next ten minutes with a digital camera placed directly in front of the test tank (Sony DCR-PJ5, Sony DCR-SX22). The videos were subsequently analyzed using the software BORIS (Friard and Gamba 2016). During the ten minutes, the number of visits (within 2 body lengths, i.e., approximately 6 cm from the patch) made by the test male towards either of the patches containing females (associate with or without predator) was noted. Additionally, total time spent in each patch, mean time spent per visit per patch and the average time difference between two successive visits to each patch were noted, to measure the association tendency of the test subject. The association preferences were measured for 27 test males.

### Experimental setup for competition experiment

The observations were conducted in a square glass arena (83× 83 cm), such that the distance from the center of the arena to any corner measured up to 40 cm, approximately ten fish standard body lengths (Figure 2). A square chamber (10×10 cm) separated by transparent perforated mesh was placed at each corner of the square arena, for housing 8 individuals (males and females), to serve as stimuli fish for the test male subject. The transparent mesh allowed visuo-chemical interactions without allowing the stimuli fish to leave their chamber. The center of the arena was provided with a removable chamber (with perforations) for acclimation of test males prior to the trials.

**Figure 2.**
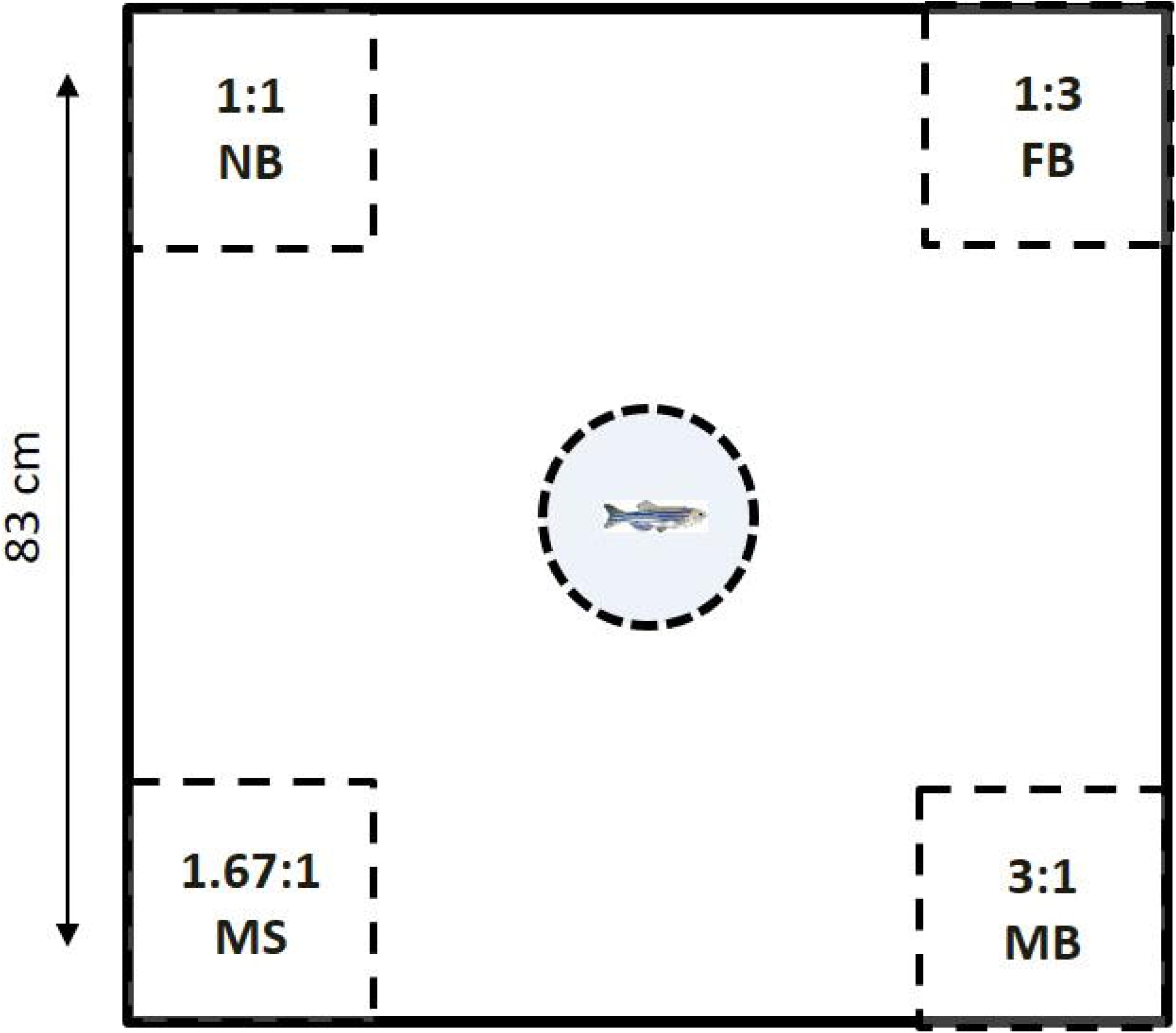
Schematic diagram of the experimental setup for testing male association preferences when presented with patches varying in male:female sex ratios. The chamber in the centre was removable and contained the test males during acclimation, which was removed, once the trials commenced. The corner square pockets contained males and females of varying sex-ratios (No Bias, Male Bias, Female Bias and Female Skewed). The corner patches were separated from the main chamber by transparent mesh partition, to allow visual and olfactory exchange.

Each corner chamber was provided with eight fishes in mixed sex ratios: four males and four females (1:1 group) (‘No Bias’ or NB group), six males and two females (3:1 group) (‘Male Bias’ or MB group), two males and six females (1:3 group) (‘Female Bias’ or FB group) and five males and three females (1.67:1 group) (‘Male Skewed’ or MS group). These chambers represented patches with differing competition levels for the test male.

Experimental protocol and subsequent video analyses was similar to the predation experiment described above. Stimuli fish were introduced into their chambers at least one hour before the commencement of the trials. At the start of trials, one test male was introduced into the acclimation chamber. After 5 min. the acclimation chamber was removed, allowing the test male to move freely in the arena. After the release of the test male, video recordings were taken for ten minutes, using a digital camera (Sony DCR-PJ5, Sony DCR-SX22) placed perpendicularly above the arena. A total of 34 subject males were tested following this protocol.

## Statistical analyses

All statistical analyses were performed in R studio (Team RC 2014: version 1.1.463). As behavioral data were not normally distributed, non-parametric paired Wilcoxon tests were performed to compare differences in patch preferences with respect to the measured parameters (i.e. total number of visits per patch; mean time spent per visit per patch and ; the average time difference between two successive visits to each patch) within each experimental condition (E1, E2, E3). For the competition experiment, we used data from the 10 minutes of behavioral video recordings of association of subject males with the patches. We constructed Generalized Linear Mixed Models (GLMMs) using package ‘lme4’ (version 1.1-19; Bates et al. 2017) in R, to build predictive models for each measured parameter, with ‘fish ID’ as the random factor and ‘Patches’ as the fixed factor, with four levels representing the four choices for the test fish. Post-hoc paired Wilcoxon test were performed to compare between the patches for each parameter. A negative binomial function was found to fit the response parameter “total number of visits” and the appropriate link function was included in the model.

## Results

### Predation experiment

We performed separate paired Wilcoxon test to compare differences in association preferences based on the measured parameters within each of our four experimental conditions. In E1, subject males visited patches with eight females (and with predator in the vicinity) significantly more (V=45, p=0.008) compared to the two-female patch (with no predator associated nearby) (Figure 3a). The males also visited the eight-female patch more often than the two-female patch indicated by a significantly lower interval time between two successive visits to the former (V=225, p=0.006) (Figure 3b). In E2, there was no clear choice by the males as we found no significant difference in any of the parameters studied (Figure 4a, 4b). In E3, males preferred the eight-female patch (without any predator in this setup) compared to the two-female patch (associated with the predator). Males also spent significantly longer time in the eight-female patch (V=276, p<<0.01) (Figure 5a) and also spent greater time per visit (V=216, p=0.02) (Figure 5b).

**Figure 3a.**
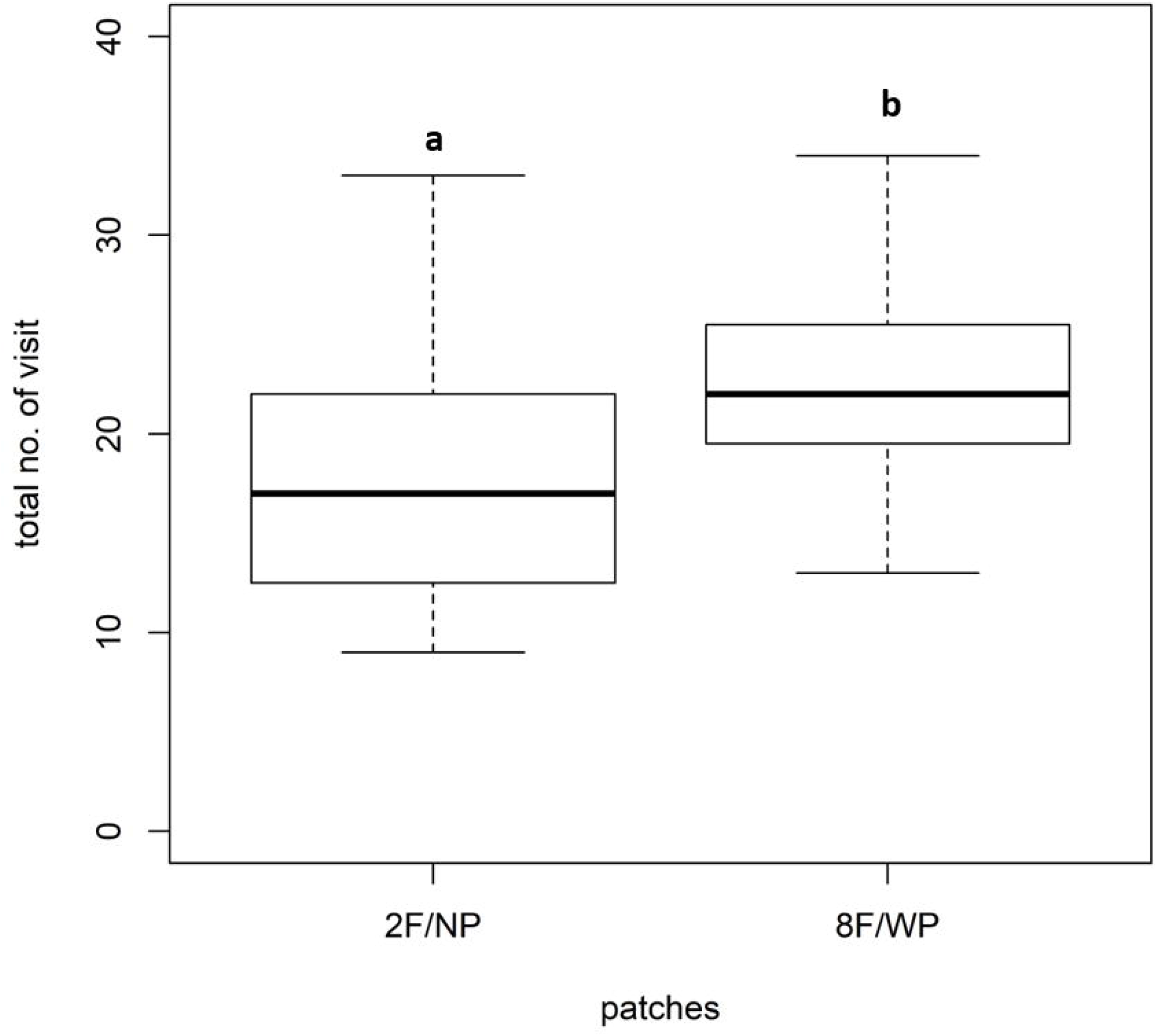
Boxplots showing the total number of visits by the male to the female-containing chambers in E1 setup of predation experiment. The chamber containing two females/no predator received significantly lesser number of visits compared to the chamber containing eight females/with predator. Similar alphabets on the top indicate no statistical difference whereas dissimilar alphabets indicate significant statistical difference (p<0.05).

**Figure 3b.**
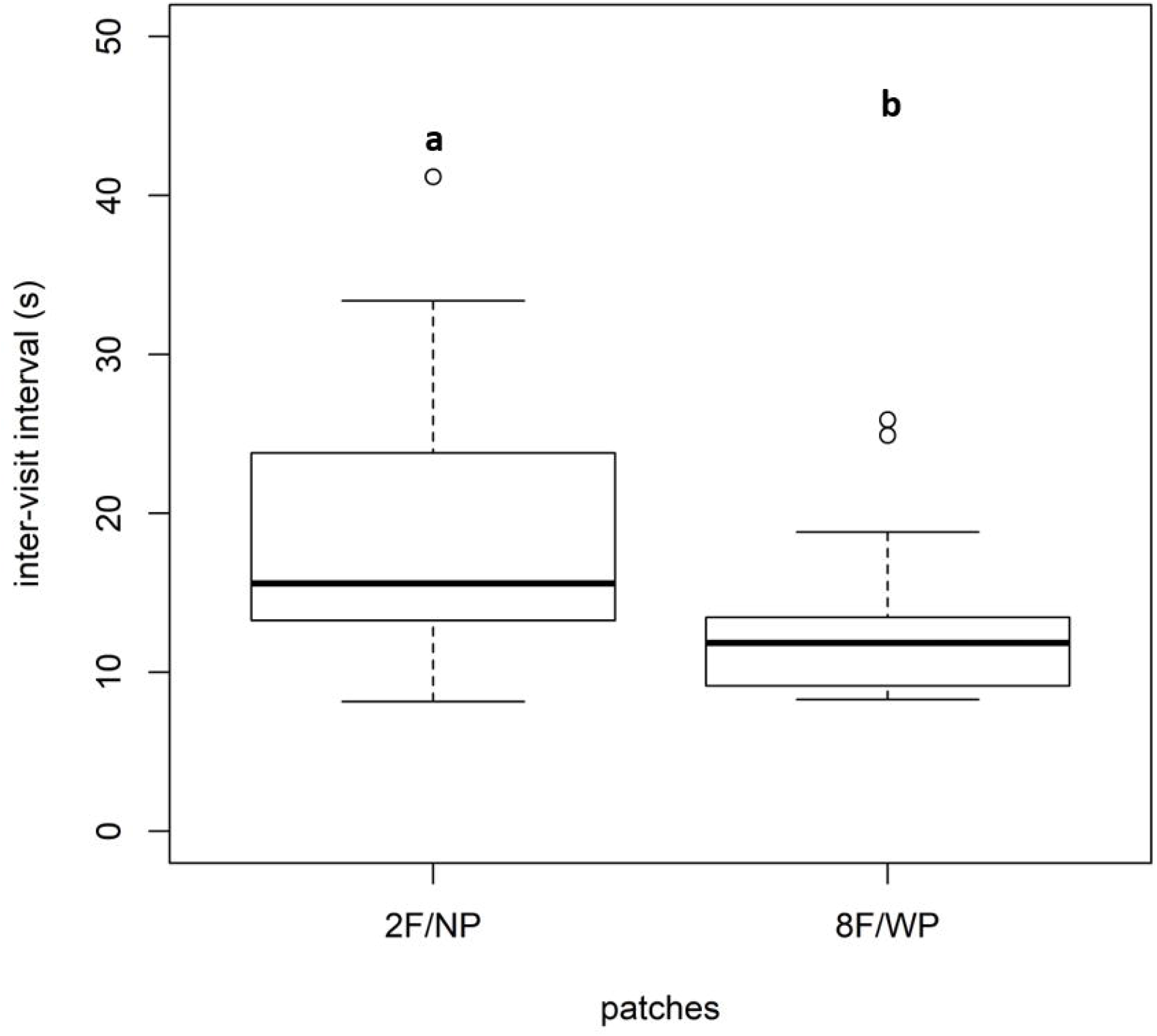
Boxplots showing the average time (in seconds) between successive visits by the male to the female-containing chambers in E1 setup of predation experiment. The chamber containing two females/no predator was visited less frequently compared to the chamber containing eight females/with predator. Similar alphabets on the top indicate no statistical difference whereas dissimilar alphabets indicate significant statistical difference (p<0.05).

**Figure 4a.**
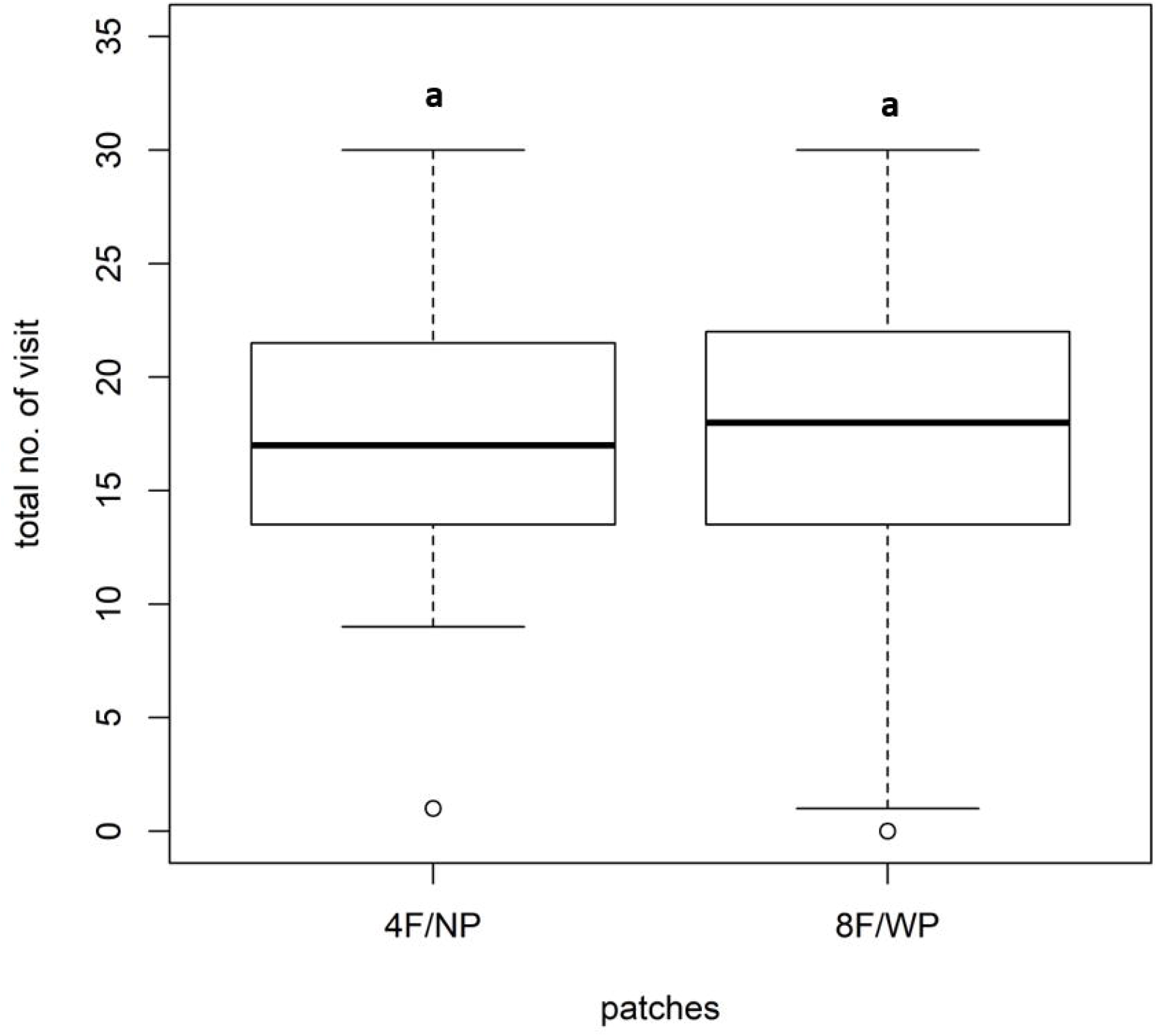
Boxplots showing the total number of visits by the male to the female-containing chambers in E2 setup of predation experiment. Both chambers received comparable number of visits by the male. Similar alphabets on the top indicate no statistical difference whereas dissimilar alphabets indicate significant statistical difference (p<0.05).

**Figure 4b.**
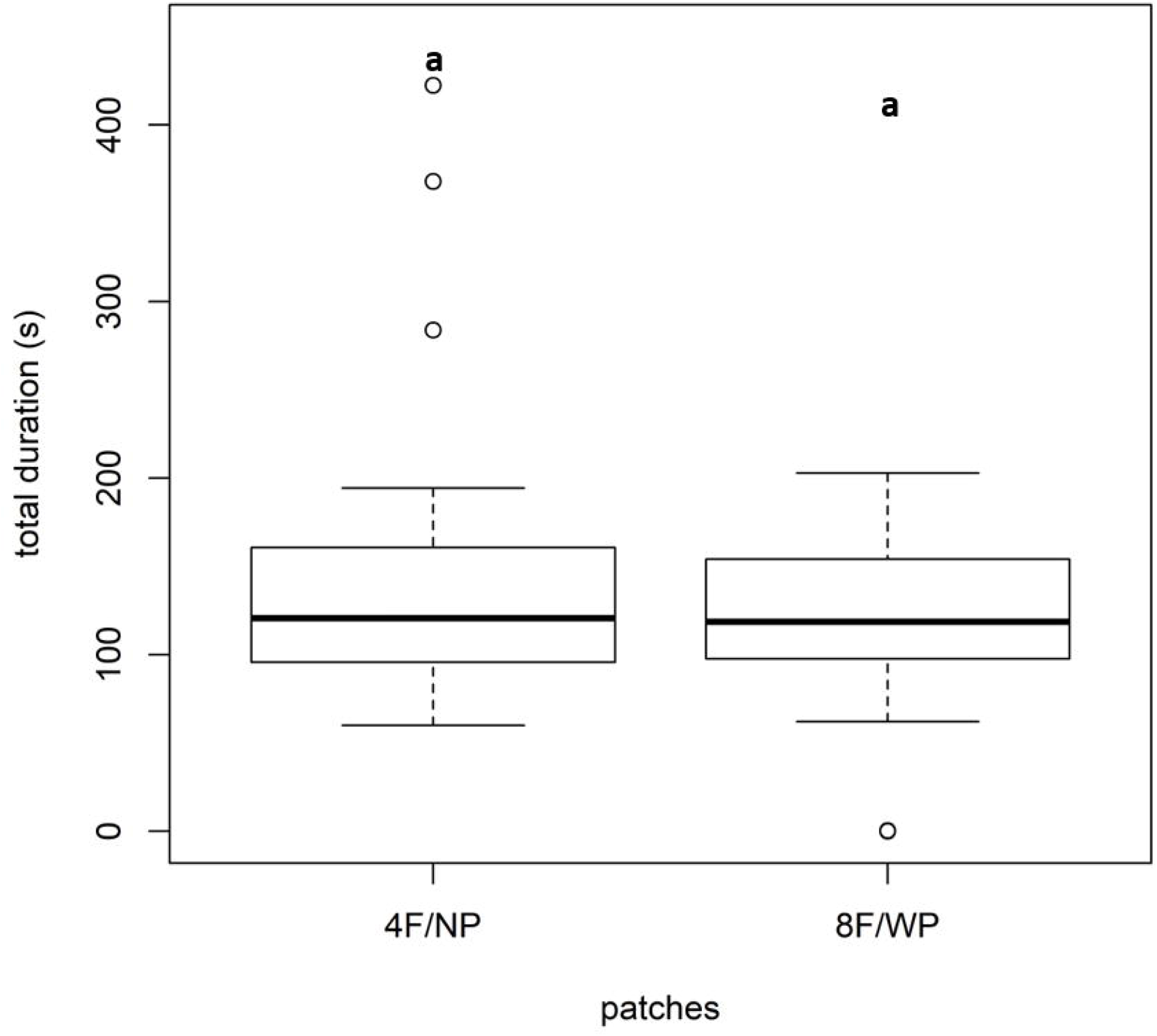
Boxplots showing the total time spent (in seconds) by the male at the female-containing chambers in E2 setup of predation experiment. Males spent comparable amount of time in both chambers. Similar alphabets indicate no statistical difference whereas dissimilar alphabets indicate significant statistical difference (p<0.05).

**Figure 5a.**
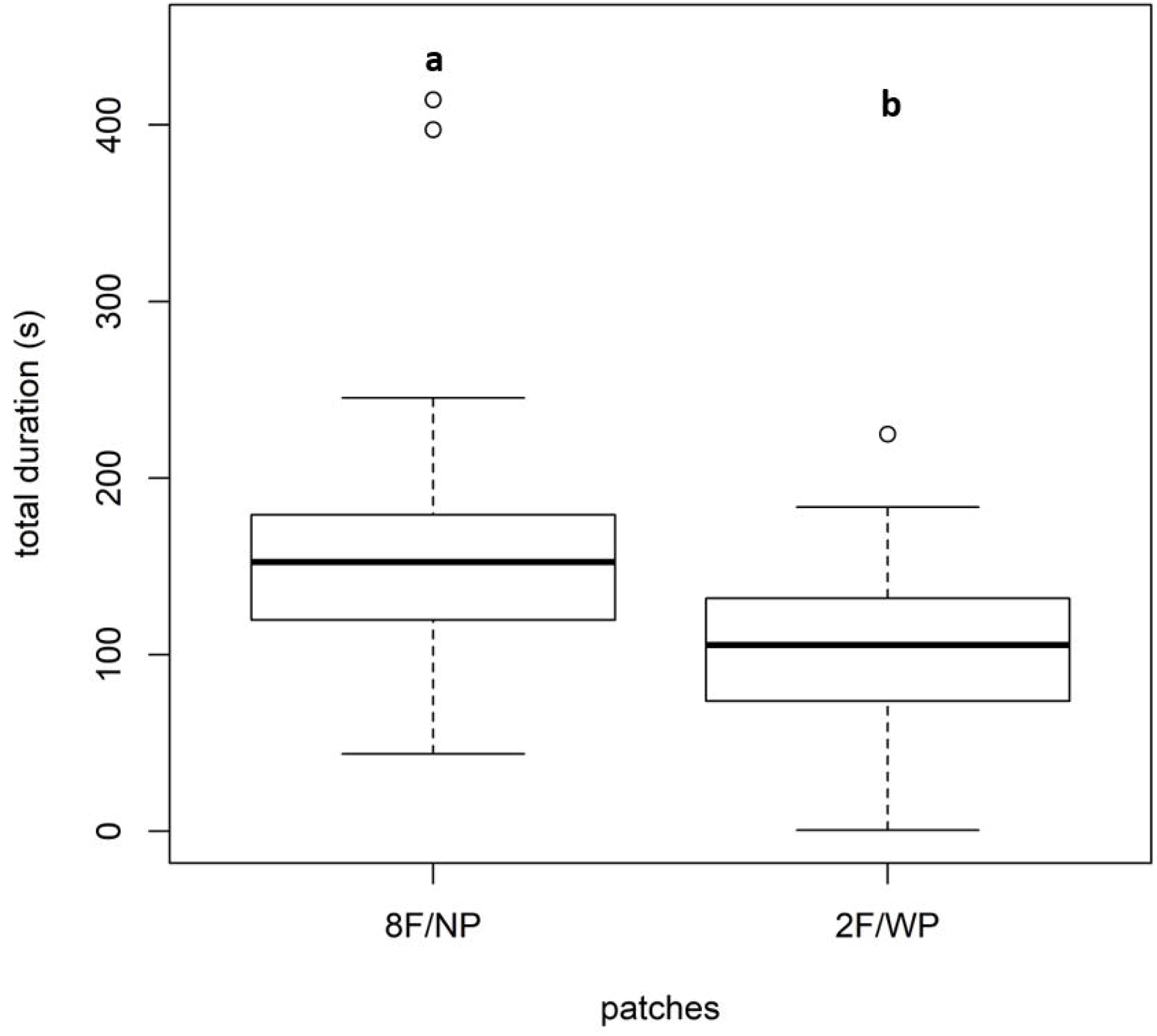
Boxplots showing the total time spent (in seconds) by the male at the female-containing chambers in E3 setup of predation experiment. Males spent greater time in eight female/no predator patch compared to the two female/without predator patch. Similar alphabets on the top indicate no statistical difference whereas dissimilar alphabets indicate significant statistical difference (p<0.05).

**Figure 5b.**
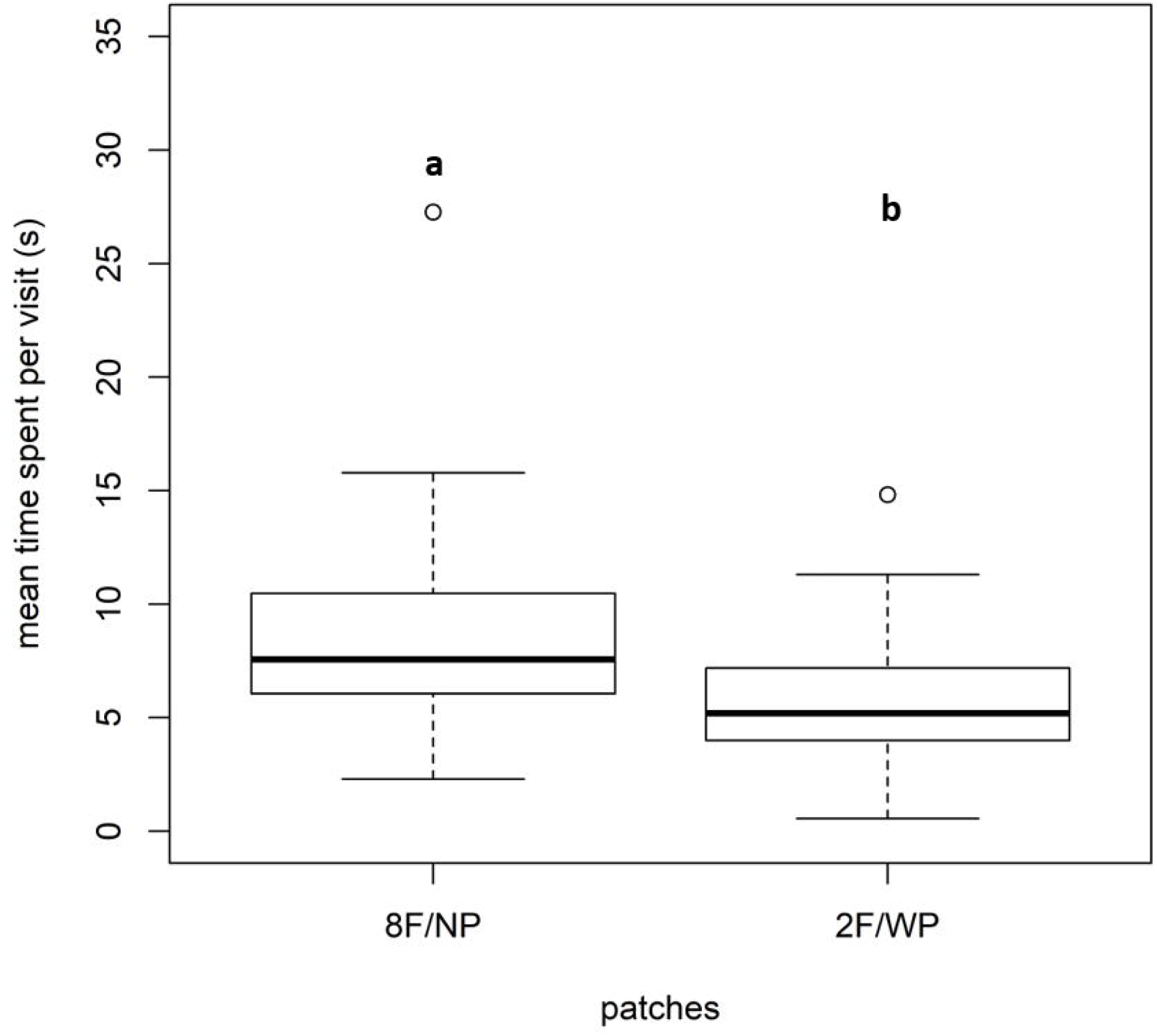
Boxplots showing the average time spent per visit (in seconds) by the male at the female-containing chambers in E3 setup of predation experiment. Males spent greater time per visit in eight female/no predator patch compared to the two female/without predator patch. Similar alphabets on the top indicate no statistical difference whereas dissimilar alphabets indicate significant statistical difference (p<0.05).

### Competition experiment

GLMM for total number of visits was found to be significantly affected by patch type (AIC=776.93, df =3, F=3.36) (Table 1a). The female biased (FB) group (6 females: 2 males) (mean=9.54sec ±1.6) received significantly lesser number of visits compared to the male skewed (MS) (5 males: 3 females) (mean=10.63sec ±1.3) (z=2.42, p=0.02) or male biased patches (MB) (6 males: 2 females) (mean=11.65sec ±1.6) (z=2.97, p=0.003) (Figure 6a). GLMM for average time spent per visit for each patch showed a significant effect of patch (AIC=933.7, df= 3, F= 3.36) (Table 1b). Significant differences were found between male biased (MB) and equal ratio (NB) patches (t=2.06, p=0.04); female biased (FB) and male biased (MB) patches (t=5.01, p<0.001); equal ratio (NB) and female biased (FB) patches (t=5.69, p<0.001) and; female biased (FB) and male skewed (MS) patches (t=-5.2, p<0.001) (Figure 6b). GLMM for total duration (AIC=1223.52, df =3, F value=1.53) (Table 1c) showed no effect of pocket and therefore, no further pair-wise comparisons were performed.

**Figure 6a.**
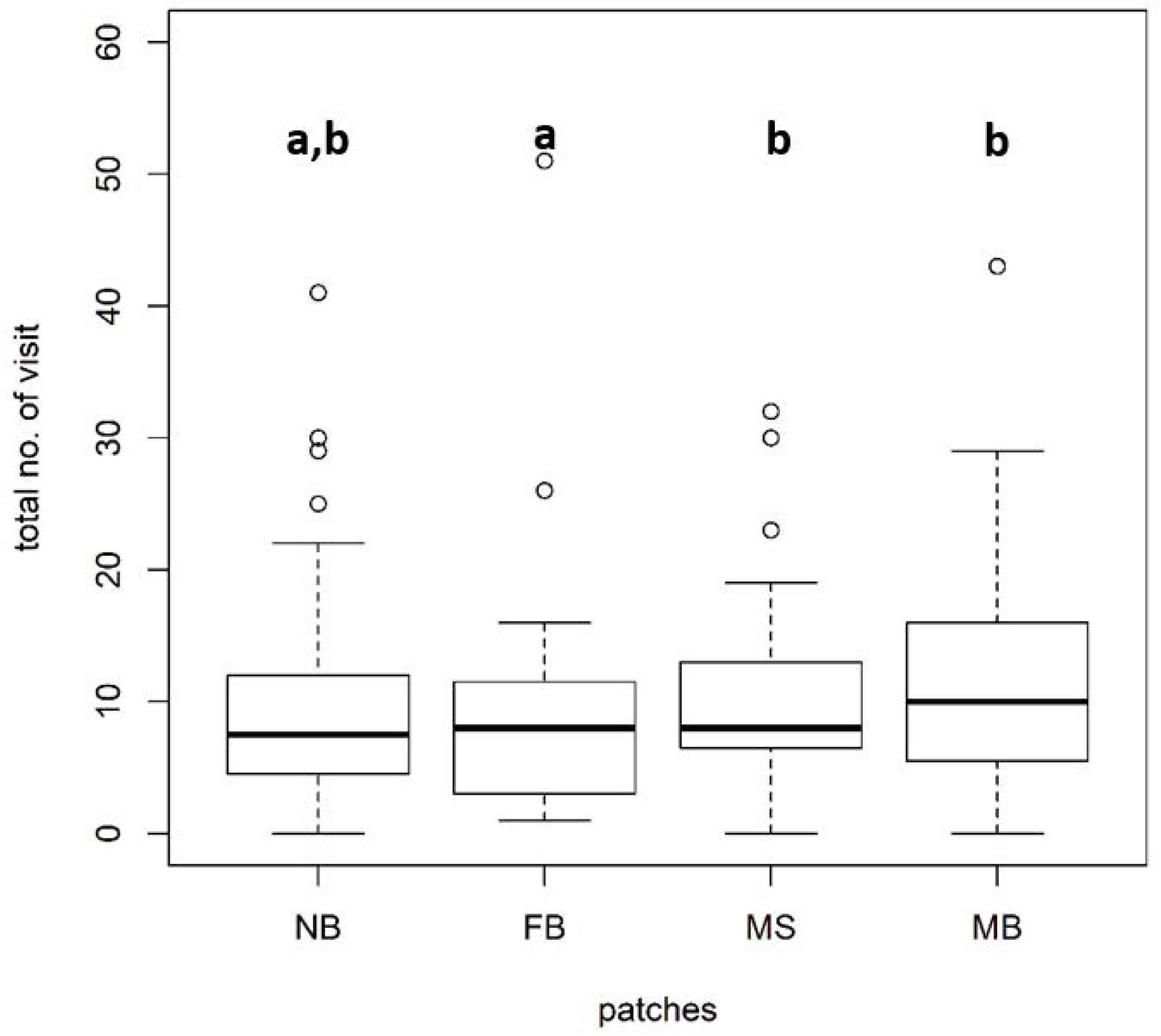
Boxplots showing the total number of visits by the male to the various chambers containing the males and females in varying sex ratios. Test males showed greater number of visits to the male biased (3:1) and male skewed patch (1.67:1) than the female-dominated patch (1:3). Similar alphabets at the top indicate no statistical difference whereas dissimilar alphabets indicate significant statistical difference (p<0.05).

**Figure 6b.**
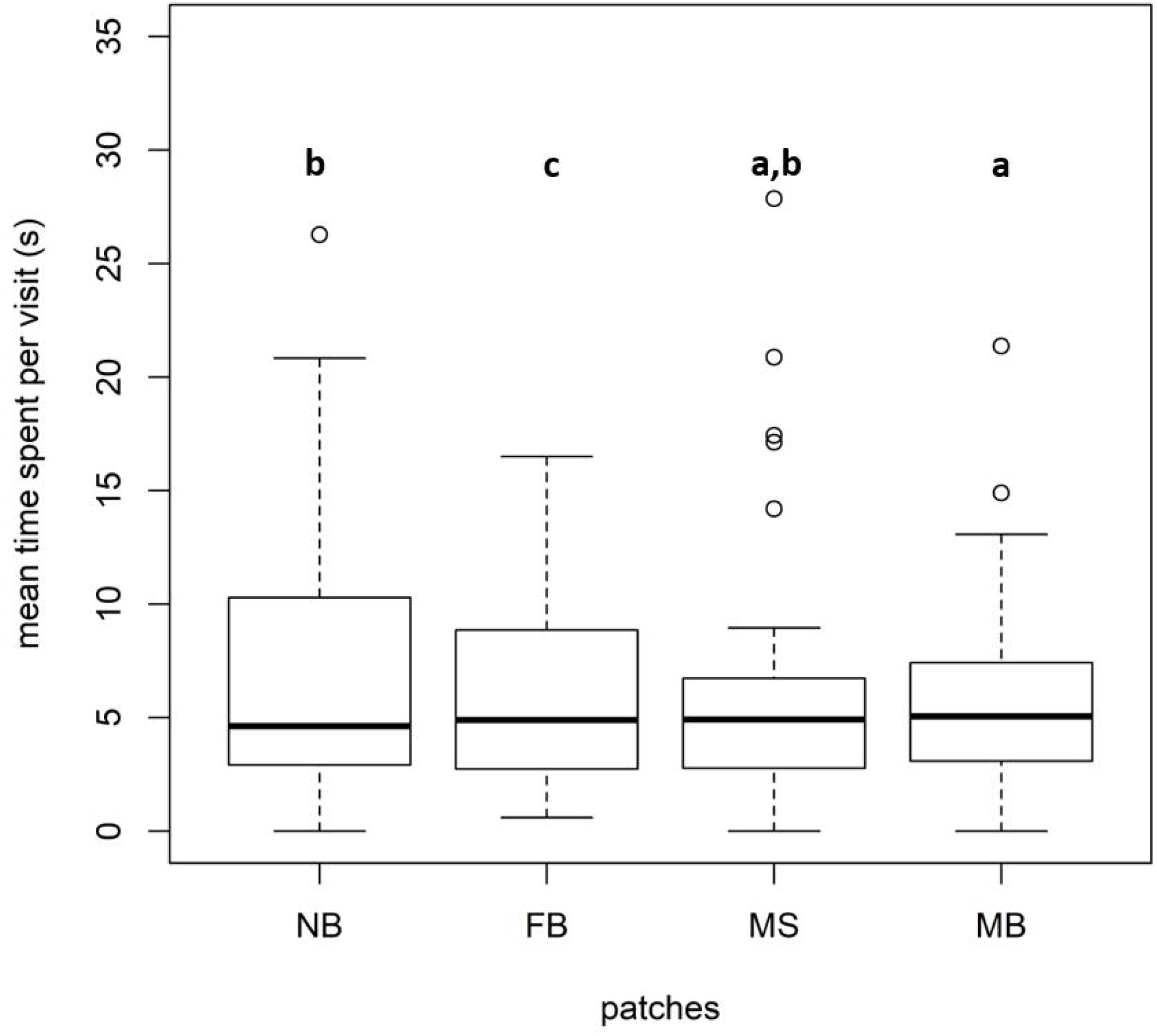
Boxplots showing the average time spent per visit by the male at the various chambers containing males and females in varying sex ratios. Test males spent greater time per patch at the female-dominated patch (1:3) compared to all other patches. Similar alphabets at the top indicate no statistical difference whereas dissimilar alphabets indicate significant statistical difference (p<0.05).

**Table 1a.**
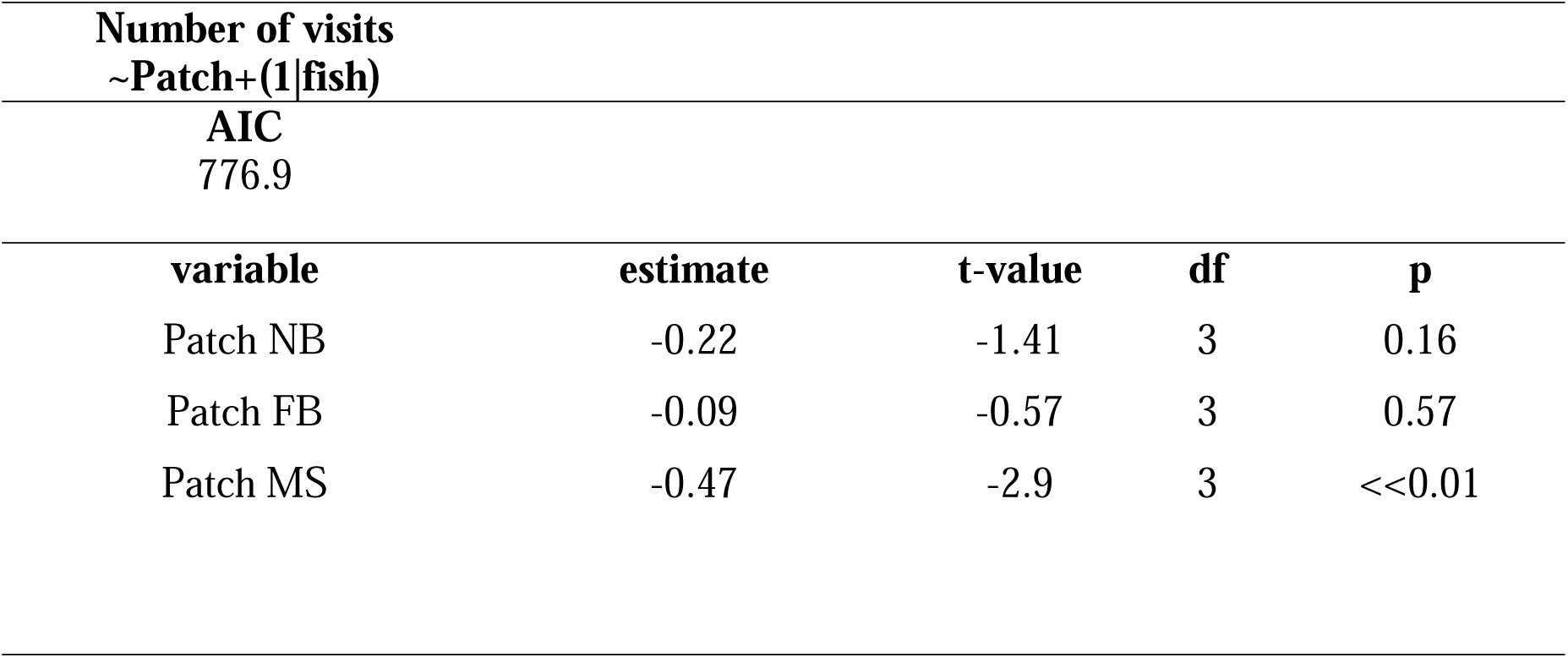
Results of the GLMM for predicting total number of visits by the male to the four mixed-sex patches, to test association preferences for patches varying in sex ratios. Model predictor estimate values, t-values, degrees of freedom (df) and p values are provided. (NB= 1:1 male: female sex ratio; FB= female biased 1:3 male: female sex ratio; MS= male skewed 1.67:1 meal: female sex ratio)

**Table 1b.** Results of the GLMM model for predicting total amount of time spent by the male to the four mixed-sex patches, to test association preferences for patches varying in sex ratios. Model predictor estimate values, t-values, degrees of freedom (df) and p values are provided. (NB= 1:1 male: female sex ratio; FB= female biased 1:3 male: female sex ratio; MS= male skewed 1.67:1 meal: female sex ratio).

|  |
| --- |
| **Total duration ~Patch+(1\|fish)** |
| **AIC** |
| 1223.5 |

| **variable** | **estimate** | **t-value** | **df** | **p** |
| --- | --- | --- | --- | --- |
| Patch NB | -0.16 | -0.73 | 3 | 0.46 |
| Patch FB | -0.04 | -0.19 | 3 | 0.85 |
| Patch MS | -0.36 | -1.7 | 3 | 0.09 |

**Table 1c.**
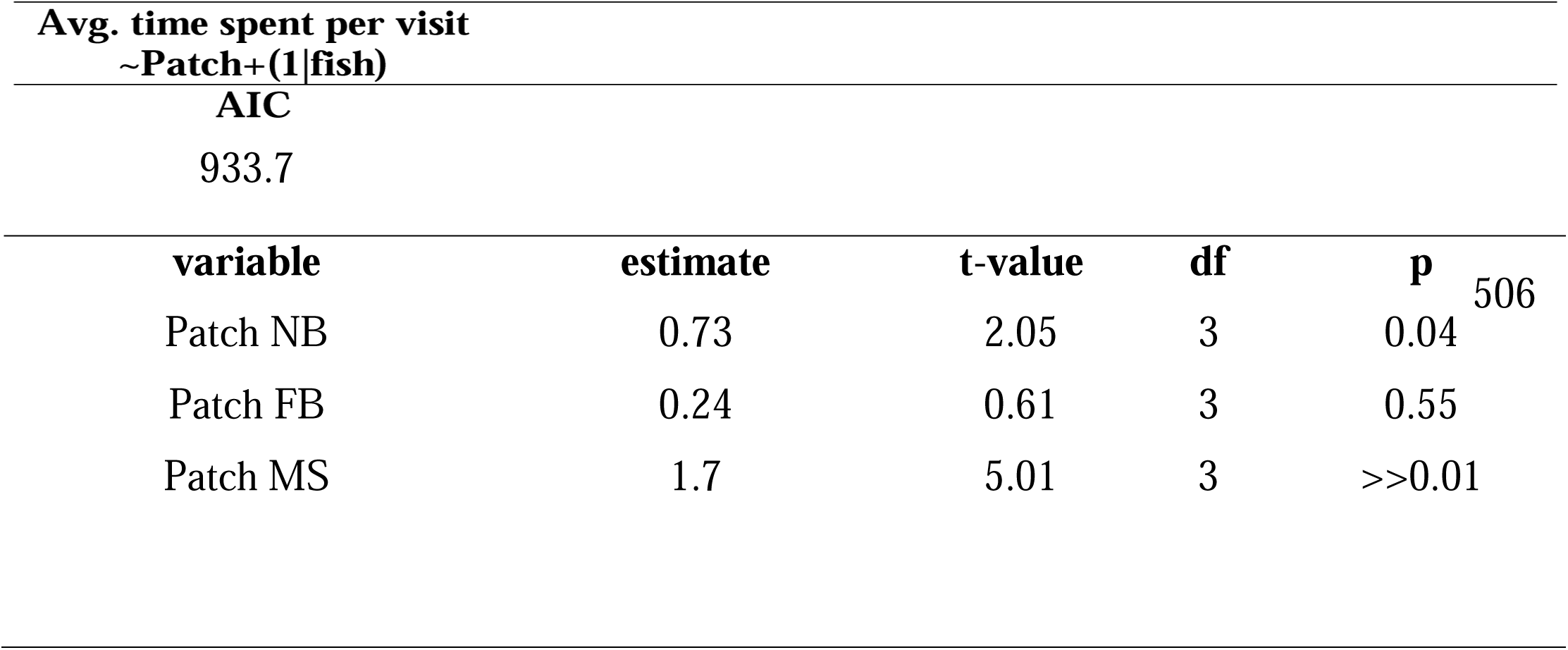
Results of the GLMM model for predicting for average time spent per visit by the male to the four mixed-sex patches, to test association preferences for patches varying in sex ratios. Model predictor estimate values, t-values, degrees of freedom (df) and p values are provided. (NB= 1:1 male: female sex ratio; FB= female biased 1:3 male: female sex ratio; MS= male skewed 1.67:1 meal: female sex ratio).

| **variable** | **estimate** | **t-value** | **df** | **p** |
| --- | --- | --- | --- | --- |
| Patch NB | 0.73 | 2.05 | 3 | 0.04 |
| Patch FB | 0.24 | 0.61 | 3 | 0.55 |
| Patch MS | 1.7 | 5.01 | 3 | >>0.01 |

## Discussion

This study explored mate searching and association preference under the influence of competition and predation threat among wild-caught zebrafish. The association choices made by subject mates when presented with patches varying in female numbers was tested under two conditions, i.e. presence of potential competitors and predator threat. Our results showed that males actually preferred to visit patches where the number of competing males were greater indicating the role of intra-sexual cues in judgment of patch quality. In the presence of predatory threat, we found that males made association decisions based on the extent of difference between the female patches associated with and without the predator.

### Association preferences for patches varying in female numbers, under predation risk

Even in the presence of a predator, males were found to show a clear preference for the eight-female patch. They visited the eight-female patch a greater number of times and also visited the former more frequently as evidenced by the shorter time interval between two successive visits. Furthermore, in the absence of a predator in the high female number patch, males showed preferential association with the eight-female patch as indicated by the higher time spent and higher average time spent per visit. Thus, irrespective of whether there is an associated risk of predator or not male zebrafish were found to prefer eight-female patch over the two-female patch, indicating that if gain in one patch is very high it is preferred over a patch with lower gain even if the former also had a higher risk.

In the condition where eight females were placed in a patch with the predator on their side and four females on the no-predator side, males did not make a clear choice for either of the female patches. Although the cost-benefit trade-off in terms of predation risk is the same for both E1 and E2 in the eight-female patch, the gains in the patch without predator is higher in E2 experimental condition compared to E1. This might account for the male making a clear choice for the eight females with predator side in E1 but not in E2.

From our results it seems likely that males do not make association decisions based on the assessment of both the patches simultaneously and not just the cost-benefit trade-off of each patch. Contrasting E1 and E2 we see increase in gains in the low risk side (no-predator side) can make the male associate more with that patch. However, when the female numbers are starkly different like in eight vs. two (in E1 and E3) the males always prefer the eight female patches irrespective of the cost associated with it. This shows a complex interaction between two ecological factors—predation pressure and number of prospective mates shaping the patch association decisions by male zebrafish.

Our results also indicate the role of numerical (counting) ability of fishes when choosing mate patches under predation pressure. They seem to easily differentiate the larger set of females when the alternative choice is much lower (e.g. 8 females vs. 2 females patch options). However, they are poorer in choosing the larger when it is 8 females vs. 4 females or 4 females vs. 2 females (see supplementary material) conditions. The ability to discriminate can have adaptive significant in organisms in the context of foraging, mating, parental care, and anti-predator strategies (Agrillo et al. 2017). Indeed, numerical abilities in fish are believed to be comparable to several mammal and bird species (Agrillo and Bisazza 2018). Guppies have been known to make clearer choices for larger numbered food patches when given a choice of one vs. four and two vs. four food items but failed to discriminate between two vs. three or three vs. four food items (Xiccato et al. 2015). Zebrafish also have been shown to clearly discriminate between greater contrasting numbers than closer ones (Potrich et al. 2015). Recent studies with angelfish and zebrafish to test their counting ability in shoal choice showed that they are able to a clear choice for larger shoals when the options are in the ratio above 2:1 but failed to make clear choices when the ratio was lesser (Gomez-Laplaza and Gerlai 2010; Seguin and Gerlai 2017). The choice of the individuals would depend on several factors such as encounter rate with predators, predator lethality and the forager’s survivor’s fitness (Brown 1999). Our results also match with studies on choice of foraging patches under predation threat observed in mule deer (Altendorf et al. 2001).

Predation pressure is an important ecological factor with survival consequences. Individuals would have to be flexible in their mate choices and modulate their search strategies based on the perceived predation risk. Our work shows that higher number of females makes the males seek a patch more frequently even when there is an immediate predation threat. However, the choice of a patch is also dependent on the other available patches and also cognitive property of the species in determining and comparing the number of prospective mates in the available patches. Thus, complex strategy involving ecological and demographic and cognitive factors in cost-benefit assessment of a patch together would be expected to determine the ultimate decision taken by individuals. Additionally, the physiological quality of females would also affect male choice and decision making. Further work is needed to understand how the quality of females determine patch association in absence and presence of predation risks for the male zebrafish.

### Association preferences for patches varying in sex ratio

Our studies reveal that total number of visits and average duration spent per visit in each patch was a function of the sex ratio of males and females comprising the patch. The test male showed significantly higher number of visits to the male-dominated (3:1) and male-skewed patch (1.67:1) than the female-dominated patch (1:3). However, average time spent per visit was significantly higher for the female-dominated patch compared to the other three patches. Thus, although they visited female-dominated patches fewer number of times, the males spent more time per visit sampling the patch. The males also spent significantly greater time per visit in the equal sex ratio patch than the male-dominated patch. Thus, there seems to be an inverse relationship between the total number of visits made to a patch and the average time spent per visit in each patch.

Our results are in contradiction to expectations that males would preferentially associate with female-dominated patches. Association preferences are likely to be driven by a combination of social as well as mating related motivations. Male preferences can also be driven by preferences and choices made by conspecific males, also referred to as ‘mate choice copying’ (MCC). It is known in several animal species that individuals choose a prospective mate based on other individuals courting that same mate (Vakirtzis 2011; Valone 2007). This can reduce sampling cost of an individual and provide honest assessment of the mates based on the assessment of the competing individuals. Female mate choice copying is also commonly observed across the animal kingdom (Pruett-Jones 1992; Godin et al. 2005; Dugatkin 1998). MCC in males among several fish species like guppies (Zimmer et al. 2014; Auld and Godin 2015), sailfin mollies (Schlupp and Ryan 1997), Atlantic mollies (Bierbach 2011) and three-spined sticklebacks (Frommen 2009) have also been documented. However, to the best of our knowledge, literature on MCC driven mate choice in zebrafish in either sexes is lacking. Further extensive studies need to be carried out to understand which of these factors, i.e., social motivation and/ choice made based on the preference of conspecific males are likely to play a role in male association.

Our work shows how zebrafish males make choices for female patches in presence of two of the most important ecological factors—predation and competition. The evaluation of female numbers in predation presence sheds light on the numerical counting cognitive ability in the species. On the other hand, preference for male-biased patches even in the presence of higher gain female-biased patches are indicative of social cues being used for patch evaluation. Together, these show a complex role of cognitive abilities and as well as social information processing in mating strategies. The current literature on zebrafish mating strategies are severely lacking and most behavioral works are based on lab-bred fishes. In contrast, our work is based on wild-caught fishes and the abundance of genomic and molecular biological tools in zebrafish can allow for deeper mechanistic studies to understand decision making involved in mate evaluation and mate choice in a sexually monomorphic species.

## Acknowledgements

The authors would like to thank the Indian Institute of Science Education and Research (IISER) - Kolkata (India) for providing infrastructural and financial support for the study. AG received Junior and Senior Research Fellowships from University Grants Commission (UGC), Government of India. Help from Rubina Mondal and Danita Daniel for routine maintenance and upkeep of laboratory and fish tanks is deeply appreciated.

## Conflict of interest

The authors declare that they have no conflict of interest.

